# Controlling the SARS-CoV-2 Spike Glycoprotein Conformation

**DOI:** 10.1101/2020.05.18.102087

**Authors:** Rory Henderson, Robert J Edwards, Katayoun Mansouri, Katarzyna Janowska, Victoria Stalls, Sophie Gobeil, Megan Kopp, Allen Hsu, Mario Borgnia, Rob Parks, Barton F. Haynes, Priyamvada Acharya

**Affiliations:** Duke Human Vaccine Institute, Durham NC 27710, USA; Duke University, Department of Medicine, Durham NC 27710, USA; Genome Integrity and Structural Biology Laboratory, NIEHS, NIH, Department of Health and Human Services, Research Triangle Park, NC 27709, USA; Duke University, Department of Immunology, Durham NC 27710, USA; Duke University, Department of Surgery, Durham NC 27710, USA

**Author notes:** To whom correspondence should be addressed Correspondence to: Rory Henderson and Priyamvada Acharya.

## Abstract

The coronavirus (CoV) viral host cell fusion spike (S) protein is the primary immunogenic target for virus neutralization and the current focus of many vaccine design efforts. The highly flexible S-protein, with its mobile domains, presents a moving target to the immune system. Here, to better understand S-protein mobility, we implemented a structure-based vector analysis of available β-CoV S-protein structures. We found that despite overall similarity in domain organization, different β-CoV strains display distinct S-protein configurations. Based on this analysis, we developed two soluble ectodomain constructs in which the highly immunogenic and mobile receptor binding domain (RBD) is locked in either the all-RBDs ‘down’ position or is induced to display a previously unobserved in SARS-CoV-2 2-RBDs ‘up’ configuration. These results demonstrate that the conformation of the S-protein can be controlled via rational design and provide a framework for the development of engineered coronavirus spike proteins for vaccine applications.

## Introduction

The ongoing global pandemic of the novel SARS-CoV-2 (SARS-2) coronavirus presents an urgent need for the development of effective preventative and treatment therapies. The viral S-protein is a prime target for such therapies owing to its critical role in the virus lifecycle. The S-protein is divided into two regions: an N-terminal S1 domain that caps the C-terminal S2 fusion domain. Binding to host receptor via the Receptor Binding Domain (RBD) in S1 is followed by proteolytic cleavage of the spike by host proteases^1^. Large conformational changes in the S-protein result in S1 shedding and exposure of the fusion machinery in S2. Class I fusion proteins, such as the CoV-2 S-protein, undergo large conformational changes during the fusion process and must, by necessity, be highly flexible and dynamic. Indeed, cryo-electron microscopy (cryo-EM) structures of SARS-2 spike reveal considerable flexibility and dynamics in the S1 domain^1,2^, especially around the RBD that exhibits two discrete conformational states – a ‘down’ state that is shielded from receptor binding, and an ‘up’ state that is receptor-accessible.

The wealth of structural information for β-CoV spike proteins, including the recently determined cryo-EM structures of the SARS-2 spike^1-11^, has provided a rich source of detailed geometric information from which to begin precise examination of the macromolecular transitions underlying triggering of this fusion machine. The transmembrane CoV S-protein spike trimer is composed of interwoven protomers that include an N-terminal receptor binding S1 domain and a C-terminal S2 domain that contains the fusion elements (Figure 1A and B).^2^ The S1 domain is subdivided into the N-terminal domain (NTD) followed by the receptor binding domain (RBD) and two structurally conserved subdomains (SD1 and SD2). Together these domains cap the S2 domain, protecting the conserved fusion machinery. Several structures of soluble ectodomain constructs that retain the complete S1 domain and the surface exposed S2 domain have been determined. These include SARS-2^1,3^, SARS^4-8^, MERS^4,9^, and other human^2,10^ and murine^11^ β-CoV spike proteins. These structures revealed remarkable conformational heterogeneity in the S-protein spikes, especially in the RBD region. Within a single protomer, the RBD could adopt a closed ‘down’ state (Figure 1A), in which the RBD covers the apical region of the S2 protein near the C-terminus of the first heptad repeat (HR1), or an open ‘up’ state in which the RBD is dissociated from the apical central axis of S2 and the NTD. Further, cryo-EM structures strongly suggest a large degree of domain flexibility in both the ‘down’ and ‘up’ states in the NTD and RBD. While these structures have provided essential information to identify the relative arrangement of these domains, the degree to which conformational heterogeneity may be altered via mutation during the natural evolution of the virus and in a vaccine immunogen design context remains to be determined.

**Figure 1.**
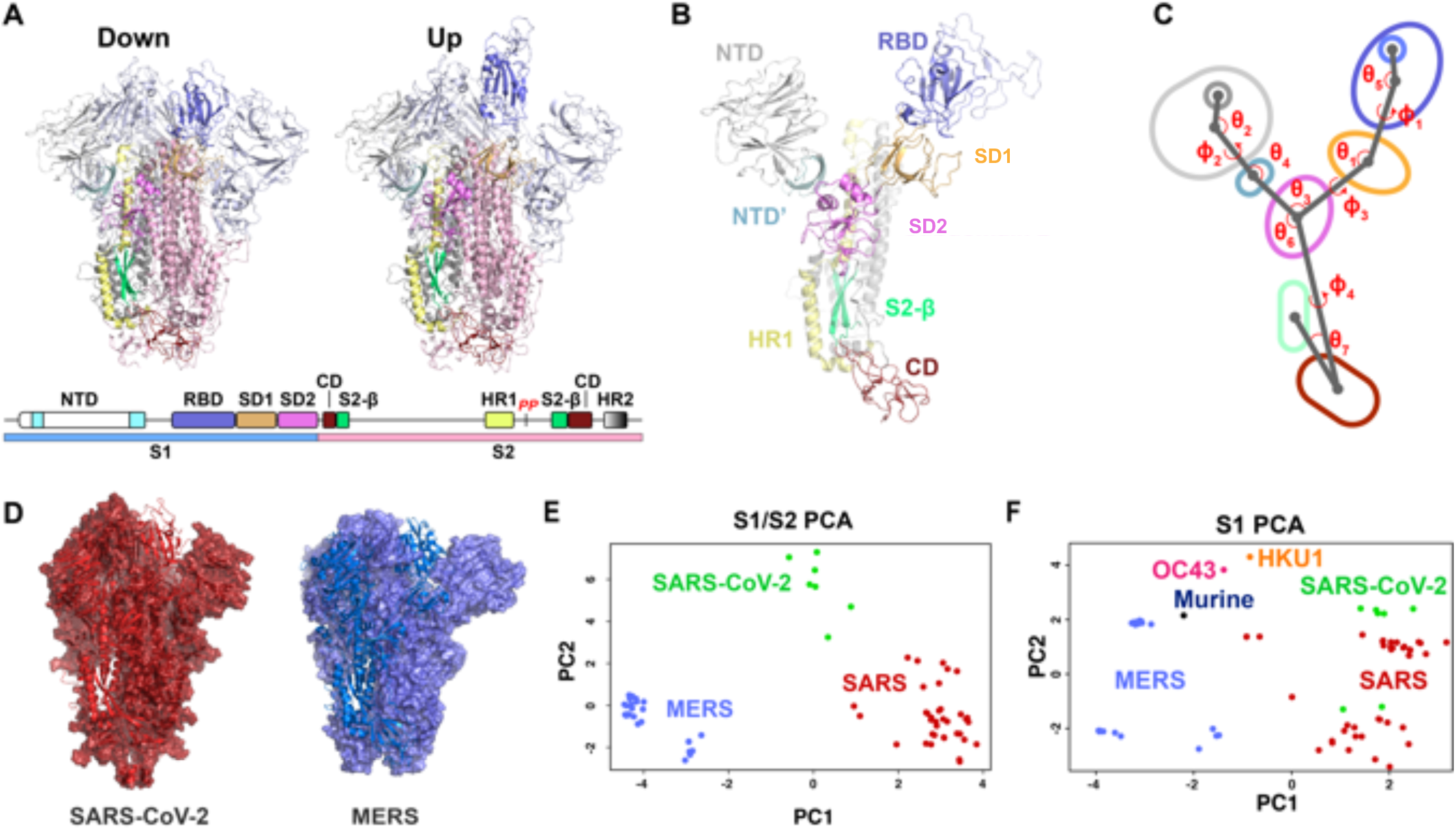
Vector based analysis of the β-CoV S-protein demonstrates remarkable variability in S-protein conformation within ‘up’ and ‘down’ states between CoV strains. **A)** Cartoon representations of the ‘down’ (upper left) and ‘up’ (upper right) state structures colored according to the specified domains (lower). **B)** A single protomer of the β-CoV S-protein with labeled domains. **C)** A simplified diagram of the β-CoV S-protein depicting the centroids and vectors connecting them with the determine angles (*****θ*****) and dihedrals (**ϕ**) labeled. **D)** The SARS-2 (left; red) and MERS (right; blue) structures each with a single protomer depicted in a cartoon representation and the remaining two in a surface representation. **E)** Principal components analysis of the SARS, SARS-2, and MERS protomers including measures between S1 and S2 domains. **F)** Principal components analysis of the SARS, SARS-2, MERS, HKU1, and Murine CoV protomers including measures only between S1 domains.

In this study we have quantified the variability in the S1 and S2 geometric arrangements to reveal important regions of flexibility to consider and to target for structure-based immunogen design. Based on these analyses, we have designed mutations that alter the conformational distribution of the domains in the S-protein. We visualized the effect of our designs using a structural determination pipeline relying first on single particle analysis by negative stain electron microscopy (NSEM) for rapid and low-cost assessment of the spike ectodomains at low resolution, followed by cryo-EM for high-resolution information on the changes introduced by these mutations. Our results reveal a heterogeneous conformational landscape of the SARS-CoV-2 spike that is highly susceptible to modification by the introduction of mutations at sites of contact between the S1 and S2 domains. We also present data on modified SARS-2 ectodomain constructs stabilized in conformations that have not yet been seen in the current available structures, with great interest and direct application in vaccine design.

## Results

### Detailed structural schema defining the geometry and internal rearrangements of movable domains of the SARS-2 spike

To characterize the unique arrangement of distinct domains in the β-CoV spike, we first aimed to develop a precise quantitative definition of their relative positions. Examination of available SARS and MERS S-protein structures revealed: 1) the NTD and RBD subdomains and internal S2 domain move as rigid bodies, and 2) these domains display a remarkable array of relative shifts between the domains in the S1 region and the S2 region’s β-sheet motif and connector domain (CD) (Figure 1B-F). In order to quantify these movements, we have analyzed the relevant regions of motion and their structural disposition in all available β-CoV ectodomain spike structures including 15 SARS^4,5,7,8^, 10 MERS^4,12^, a HKU1^2,10^, an OC43^2,10^, a murine β-CoV, and three SARS-2^13,14^ structures (Figures 1E-F and 2). Each protomer in those structures displaying asymmetric ‘up’/down’ RBD states was examined independently yielding a dataset of 76 structural states. The NTD was split into a primary N-terminal section and a secondary C-terminal section based upon visual inspection of this region in the various β-CoV structures (Figures 1B-C and 2). We next analyzed S-protein geometry using a vector-based approach. Specifically, vectors connecting each region’s C_α_ centroids were generated and used to define the relative dispositions of the domains (Figure 1C and 2). The vector magnitudes and select angles and dihedrals were used to identify the breadth of differences in domain positioning and compare between strains. The results indicated that β-CoV spike proteins in various strains differ markedly from one another and that considerable variability in the domain arrangements within strains exists, especially in the SARS ectodomains (Figures 1E-F and 2A-H). In particular, both ***θ***_1_ and ϕ_1_ (Figure 2A-B), describing the angle between the SD2 to SD1 and SD1 to RBD vectors as well as the SD1 to RBD dihedral, respectively, effectively report on the ‘up’ and ‘down’ configurations while indicating substantial differences between SARS and MERS in both the ‘up’ and ‘down’ states. The angular disposition of the NTD elements further indicated differences in SARS and MERS with a particularly marked shift from the examined β-CoV spikes in the murine structure (Figure 1E). Additional S1 differences are observed between vectors involving SD2. The disposition of the S2 domain relative to S1 defined by the dihedral about the vector connecting SD2 to the S2 CD differs markedly between MERS/SARS-2 and SARS as well with the angle between the vectors connecting the NTD’ to SD2 and SD2 to the CD demonstrating a shift in SARS-2. Finally, the disposition of the CD to the inner portion of S2 measured as an angle between a vector connected to an interior S2 β-sheet motif and the vector connecting the CD to SD2 indicates SARS differs from both MERS and SARS-2. Interestingly, the MERS disposition appears to respond to RBD triggering, displaying a bimodal distribution. These results demonstrate that, while the individual domain architectures and overall arrangements are conserved (Figure 1D), important differences between these domains exists between strains, suggesting that subtle differences in inter-domain contacts could play a major role in determining these distributions and thereby alter surface antigenicity and the propensity of the domains to access ‘up’ and ‘down’ RBD states.

**Figure 2.**
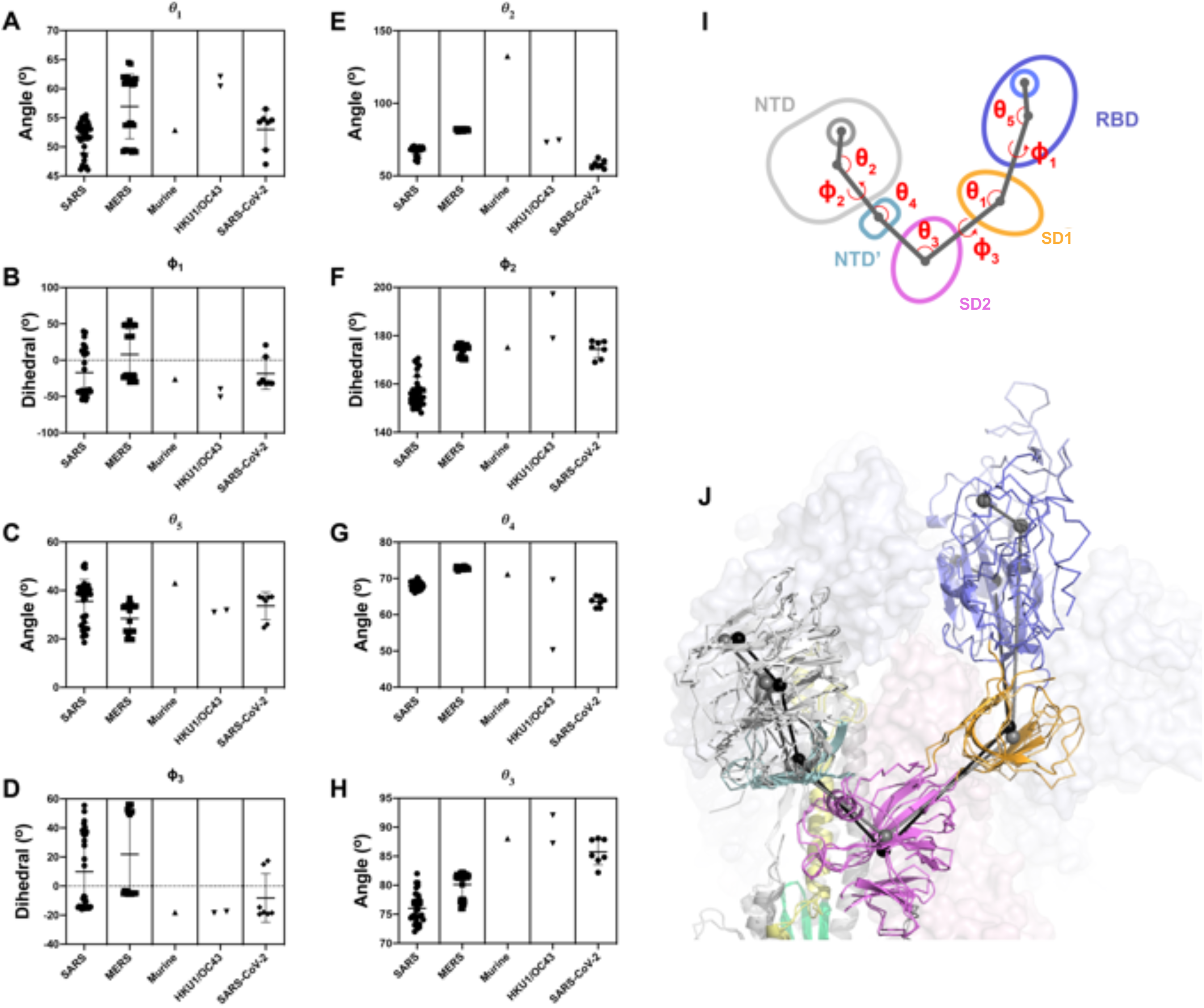
Vector based analysis of the CoV S-protein demonstrates remarkable variability in S-protein conformation within ‘up’ and ‘down’ states between CoV strains. **A)** Angle between the subdomain 1 to subdomain 2 vector and the subdomain 1 to RBD vector. **B)** Dihedral about the subdomain 1 to RBD vector. **C)** Angle between the RBD to subdomain 1 vector and the RBD to RBD helix vector. **D)** Dihedral about the subdomain 2 to subdomain 1 vector. **E)** Angle between the NTD’ to NTD vector and the NTD to NTD sheet motif vector. **F)** Dihedral about the NTD to NTD’ vector. **G)** Angle between the NTD’ to subdomain 2 vector and the NTD’ to NTD vector. **H)** Angle between the subdomain 2 to NTD’ vector and the subdomain 2 to subdomain 1 vector. **I)** Diagram of the domains and relevant angles and dihedrals for S1 **J)** Cartoon representation of one protomer’s S1 domains in the ‘down’ state overlaid with a ribbon representation of the ‘up’ state colored according to (I). Black (‘down’ state) and grey (‘up’ state) spheres represent domain centroids with lines connecting representing the vectors. Adjacent protomers represented as transparent surfaces.

### Identification of sites for differential stabilization of the SARS-2 ectodomain spike RBD orientation

Based on the observed variability in the geometric analysis of β-CoV spikes, we asked whether the propensity for the RBD to display the ‘down’ and ‘up’ states could be modified via mutations without altering exposed antigenic surfaces. To this end, we identified protomer to protomer interactive sites amenable to modification and down selected mutations at these sites using the Schrödinger Biologics suite^38,39,40^. In an effort to eliminate exposure of the receptor binding site of the RBD, we examined the potential for disulfide linkages between the RBD and its contact with S2 near the C-terminus of HR1 to prevent RBD exposure. We identified a double cysteine mutant, S383C and D985C (<u>R</u>BD to <u>S2</u> double mutant; rS2d; Supplemental Figure S1), as a candidate for achieving this goal. The transition from the ‘down’ state to the ‘up’ state involves shifts in the RBD to NTD contacts. Therefore, in an effort to prevent these shifts, we identified a site in an RBD groove adjacent to the NTD for which we prepared a triple mutant, D398L, S514L, and E516L (<u>R</u>BD to <u>N</u>TD <u>t</u>riple mutant; rNt, Supplemental Figure S1). As SD1 acts as a hinge point for the RBD ‘up’/’down’ transitions (Figures 1A-C, 2I-J), we hypothesized that enhanced hydrophobicity at the SD1 to S2 interface might shift the position of SD1, thus influencing the hinge and potentially the propensity for RBD triggering. A double mutant, N866I and A570L (<u>S</u>ubdomain <u>1</u> to <u>S2</u> <u>d</u>ouble mutant; u1S2d, Supplemental Figure S1), as well as quadruple mutant, A570L, T572I, F855Y, and N856I (S<u>u</u>bdomain <u>1</u> to <u>S2</u> <u>q</u>uadruple mutant; u1S2q), were identified for this purpose. Finally, we asked whether linking SD2 to S2 would alter the conformational distribution of the RBDs. The double cysteine mutant, G669C and T866C (S<u>u</u>bdomain <u>1</u> to <u>S2</u> <u>d</u>ouble mutant; u2S2d, Supplemental Figure S1), was identified for this purpose. These mutants were prepared in the context of a previously published SARS-2 ectodomain construct^3^.

### NSEM analysis of the SARS-2 spike ectodomain proteins

To assess the quality of the purified spike proteins and to obtain low resolution visualization of the structures, we performed NSEM analysis. The micrographs showed a reasonably uniform distribution of particles consistent with the size and shape of the SARS-2 spike ectodomain (Figure 3). 2D class averages showed spike populations with well resolved domain features. The data were subjected to 3D-classification followed by homogeneous refinement. The unmutated construct was resolved into two classes of roughly equal proportions. The two classes differed in the position of their RBD domains. One class displayed all three RBDs in their ‘down’ positions, whereas, the other class displayed one RBD in the ‘up’ position. This was consistent with published cryo-EM results^15^ that described a 1:1 ratio between the ‘down’ and ‘1-up’ states of the SARS-2 spike ectodomain. The mutant spikes were analyzed using a similar workflow as the unmutated spike. All of the mutants displayed well-formed spikes in the micrographs, as well as in the 2D class averages. Following 3D classification, for the rS2d construct, we observed only the ‘down’ conformation; the 1-RBD ‘up’ state that was seen for the unmutated spike was not found in this dataset. The u1S2q mutant presented another striking finding, where we observed a new conformational state with 2 RBDs in the ‘up’ position. The 2-RBD ‘up’ state has been reported before for the MERS CoV spike ectodomain^12^ but has not been observed thus far for either the SARS or the SARS-2 spikes. Based on the NSEM analysis we selected the rS2d and u1S2q constructs for high resolution analysis by cryo-EM.

**Figure 3.**
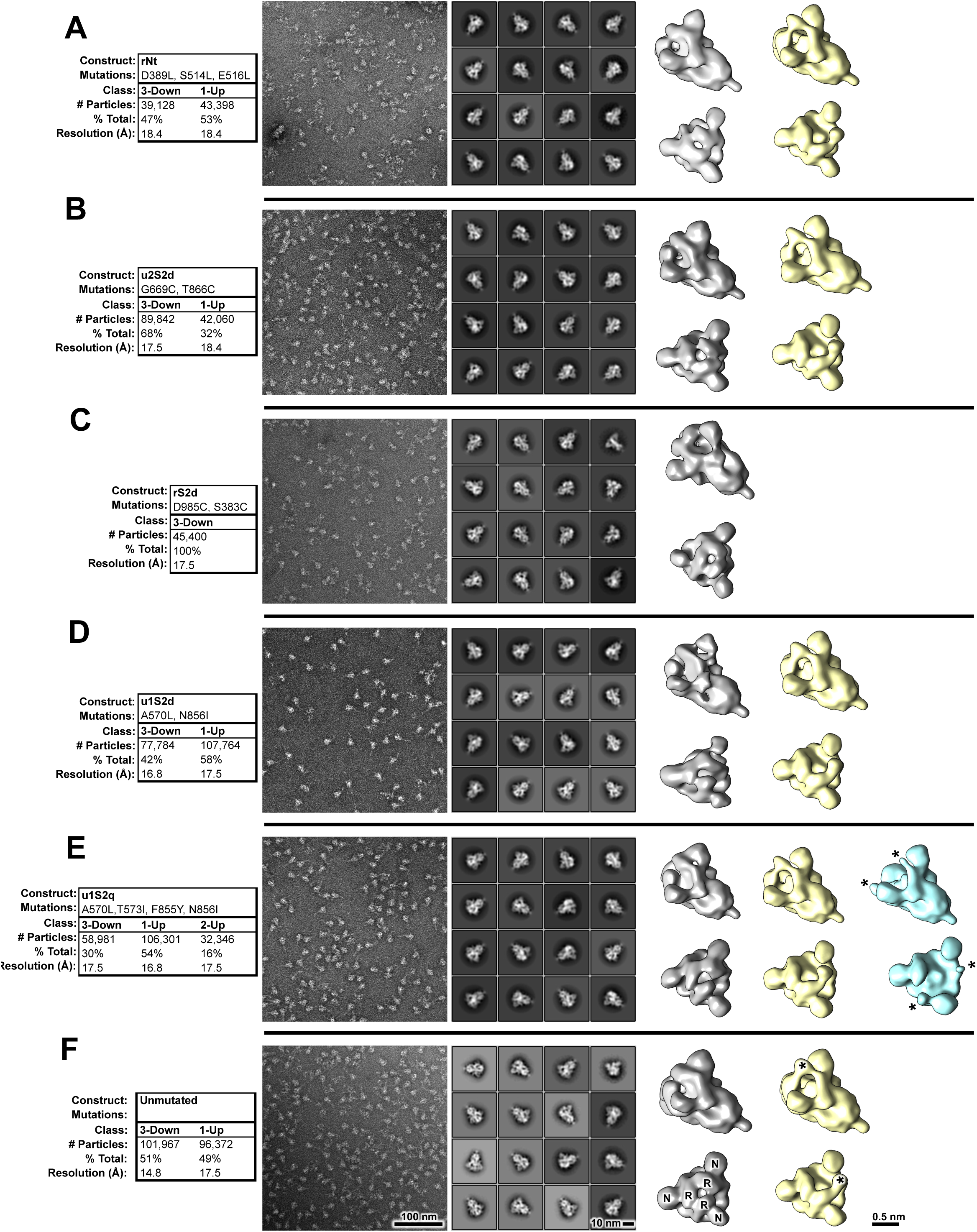
Negative stain electron microscopy analysis of S-protein constructs. **A)** Data tables, indicating construct names, mutations, observed classes, number and percent of particles per class and final resolution (gold-standard Fourier-shell correlation, 0.143 level). **B)** Raw micrographs. **C)** Representative 2D class averages. **D)** 3D reconstructions of 3-RBD-down classes, shown in top view, looking down the S-protein 3-fold axis on the left and tilted view on the right. **E)** 3D reconstructions of 1-RBD-up classes. Up-RBD is marked with an asterisk. **F)** 3D reconstruction of 2-RBD-up class. Density for up-RBDs is weak, indicated by asterisks. Receptor binding domains and N-terminal domains of first structure marked with R and N, respectively.

### Cryo-EM analysis of the SARS-2 spike ectodomain proteins

To visualize the mutations and their effect on the structure of the spike, we collected cryo-EM datasets for the rS2d and u1S2q constructs (Figure 4-7, Table 1, Supplemental Figures S2 and S3). Consistent with what was observed in the NSEM analysis, after multiple rounds of 2D and 3D-classification to remove junk particles and broken and/or misfolded spikes, we found a population of ‘down’ state spike in the rS2d dataset through *ab initio* classification in cryoSparc. We then implemented additional exhaustive *ab initio* classifications, as well as heterogeneous classifications using low-pass filtered maps of known open conformations of CoV spikes to search for open state spikes in the dataset. We were unable to find any such states, confirming that the SARS-2 spike was locked in its ‘down’ conformation in the rS2d mutant. The rS2d disulfide-linked density at the mutation site confirmed disulfide formation in the double mutant (rS2d) (Figure 5). Comparison of the domain arrangements of this construct with that of the unmutated ‘down’ closed state structure indicated the protein structure was otherwise unperturbed (Supplemental Figure S4A).

**Table 1:**
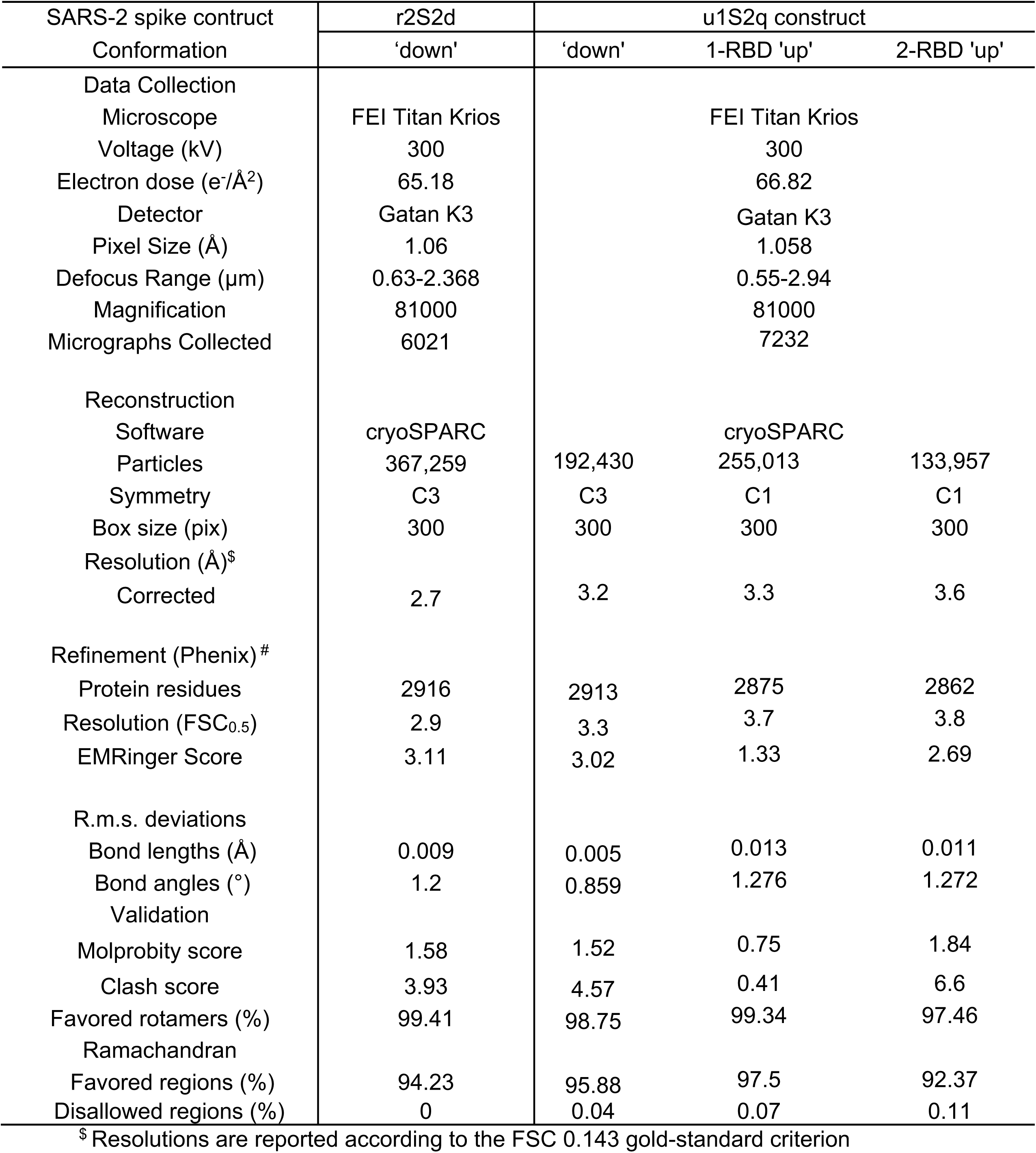
Cryo-EM Data Collection and Refinement Statistics.

| SARS-2 spike construct<br>Conformation | r2S2d | u1S2q construct |  |  |
| --- | --- | --- | --- | --- |
|  | 'down' | 'down' | 1-RBD 'up' | 2-RBD 'up' |
| Data Collection |  |  |  |  |
| Microscope | FEI Titan Krios |  | FEI Titan Krios |  |
| Voltage (kV) | 300 |  | 300 |  |
| Electron dose (e<sup>-</sup>/Å<sup>2</sup>) | 65.18 |  | 66.82 |  |
| Detector | Gatan K3 |  | Gatan K3 |  |
| Pixel Size (Å) | 1.06 |  | 1.058 |  |
| Defocus Range (µm) | 0.63-2.368 |  | 0.55-2.94 |  |
| Magnification | 81000 |  | 81000 |  |
| Micrographs Collected | 6021 |  | 7232 |  |
| Reconstruction |  |  |  |  |
| Software | cryoSPARC |  | cryoSPARC |  |
| Particles | 367,259 | 192,430 | 255,013 | 133,957 |
| Symmetry | C3 | C3 | C1 | C1 |
| Box size (pix) | 300 | 300 | 300 | 300 |
| Resolution (Å)<sup>§</sup> |  |  |  |  |
| Corrected | 2.7 | 3.2 | 3.3 | 3.6 |
| Refinement (Phenix)<sup>#</sup> |  |  |  |  |
| Protein residues | 2916 | 2913 | 2875 | 2862 |
| Resolution (FSC<sub>0.5</sub>) | 2.9 | 3.3 | 3.7 | 3.8 |
| EMRinger Score | 3.11 | 3.02 | 1.33 | 2.69 |
| R.m.s. deviations |  |  |  |  |
| Bond lengths (Å) | 0.009 | 0.005 | 0.013 | 0.011 |
| Bond angles (°) | 1.2 | 0.859 | 1.276 | 1.272 |
| Validation |  |  |  |  |
| Molprobity score | 1.58 | 1.52 | 0.75 | 1.84 |
| Clash score | 3.93 | 4.57 | 0.41 | 6.6 |
| Favored rotamers (%) | 99.41 | 98.75 | 99.34 | 97.46 |
| Ramachandran |  |  |  |  |
| Favored regions (%) | 94.23 | 95.88 | 97.5 | 92.37 |
| Disallowed regions (%) | 0 | 0.04 | 0.07 | 0.11 |

<sup>§</sup> Resolutions are reported according to the FSC 0.143 gold-standard criterion

**Figure 4.**
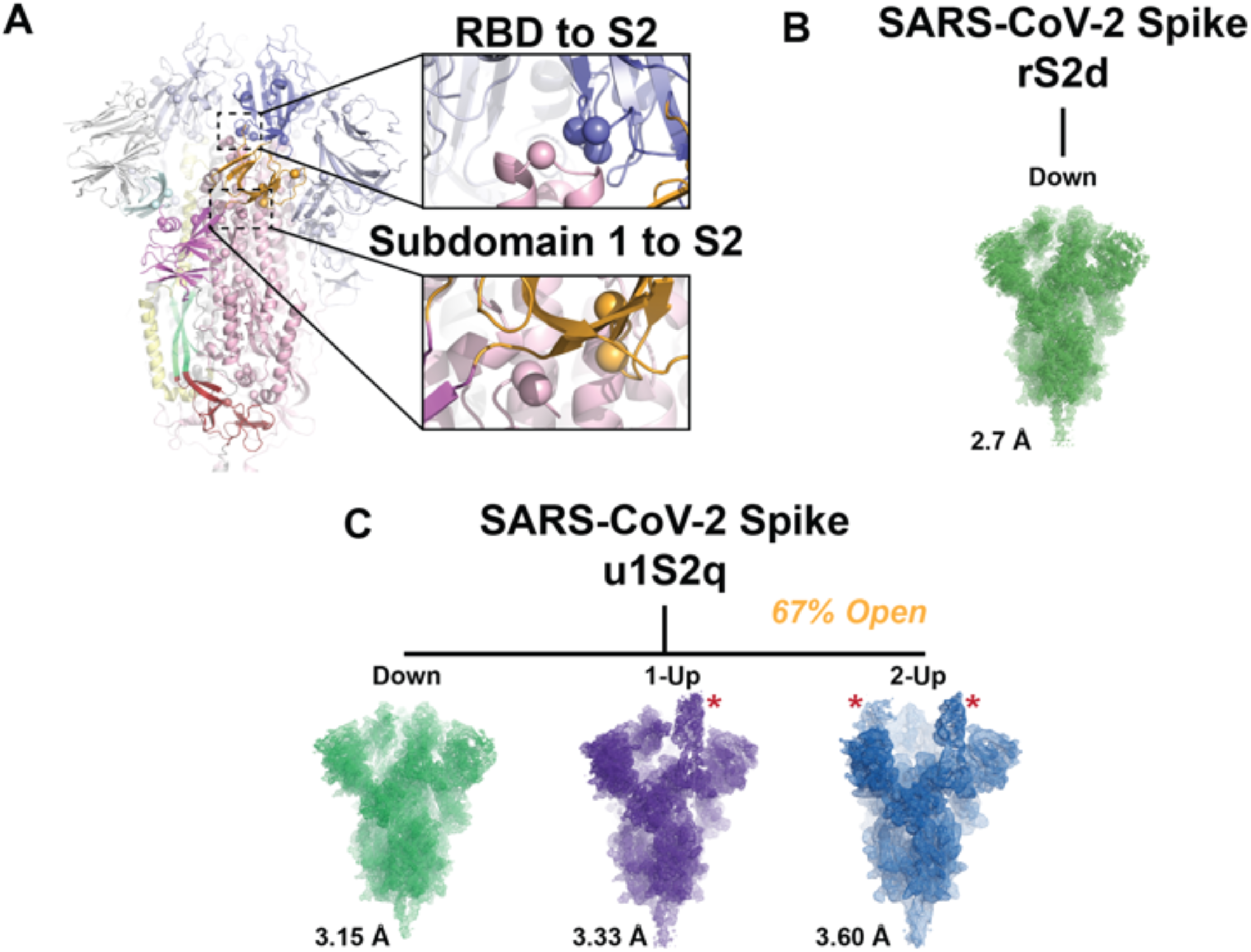
Cryo-EM dataset reveals differential stabilization of the S-protein in the mutant ectodomain constructs. **A)** The S-protein spike highlighting the two regions of interest for structure and computation-based design. **B)** (*top*) The rS2d RBD to S2 locked structure displaying only the all RBD down state. (*bottom*) **C)** (*top*) The u1S2q SD1 to S2 mutated structure displaying the all RBD ‘down’ state, the 1-RBD ‘up’ state, and, for the first time in the SARS-2 S ectodomain, the 2-RBD ‘up’ state.

**Figure 5.**
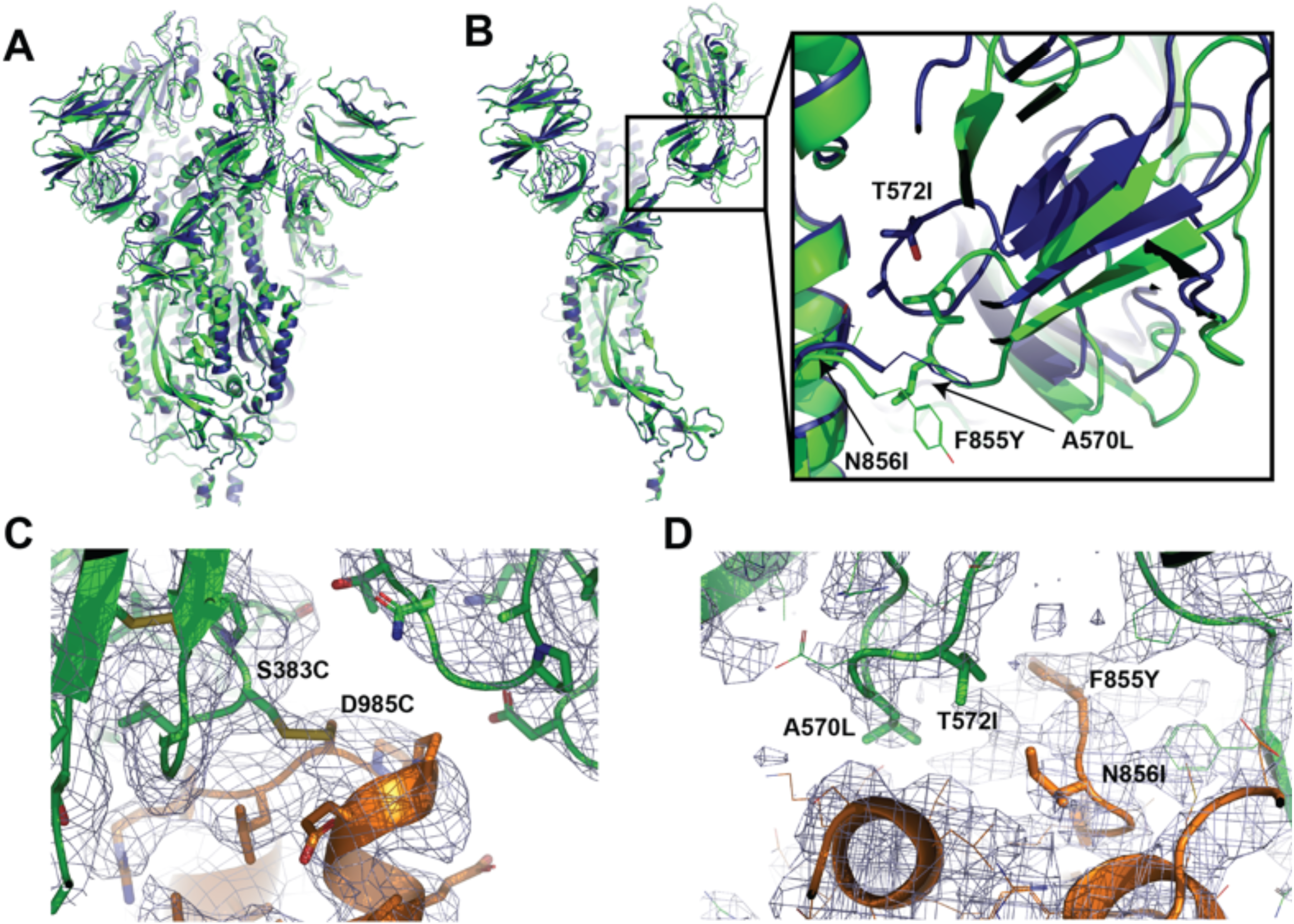
Cryo-EM structures of the “down” state in the r2S2d and u1S2q constructs reveal differential stabilization of domain positions. **A)** Alignment between the trimers of the designed disulfide linked rS2d (dark blue) mutant structure and the u1S2q (green). **B)** (left) Alignment between single protomers of the designed disulfide linked rS2d(dark blue) mutant structure and the u1S2q(green). (right) Zoomed in view of SD1 in both constructs demonstrating the shift in the subdomain with the 4 mutants. **C)** Structure and cryo-EM map depicting the RBD to S2 bridging density between the introduced cysteine residues. **D)** Structure and cryo-EM map depicting the SD1 and S2 mutations.

In contrast to the rS2d design, the u1S2q design displayed widespread rearrangement of the S1 domains (Supplemental Figure S1B). In the ‘down’ state structure, density in the mutated S2 position remained in the configuration observed in the unmutated construct, with the N855I and F856Y residue loop in close proximity the S2 residue L966 and S1 residue P589. This suggested these mutations had little impact on the observed shifts. However, the S2-interactive SD1 displayed a rigid body movement relative to both the rS2d and unmutated constructs with ***θ***_1_ and ϕ_3_ displacements of 3.4° and 1.8°, respectively (Figure 5A and B). This resulted in displacement of the A570L+T572I containing loop from the unmutated position which resides near the S2 L966 residue (Figure 5B and Supplemental Figure S1C). The S2 contact disruption is accompanied by an angular shift of the NTD away from the primary trimer axis owing to SD1 to NTD’ contacts, yielding ***θ***_3_ and ϕ_3_ shifts of 5.4° and 7.7°, respectively (Figure 6C). The subdomain rearrangement impacts the positioning of the RBD with only a minor shift in the ϕ_1_ dihedral of 0.1° indicating the RBD moved with SD1 indicated in the ***θ***_1_/ϕ_3_ shifts. The newly acquired arrangement in both the RBD and NTD was further accompanied by an apparent increase in their flexibility suggesting conformational heterogeneity. These ‘down’ state shifts were observed in both the single RBD ‘up’ structure and the two RBD ‘up’ structures (Figure 6). Interestingly, the extent to which the SD1 shift differed from that observed in the unmutated construct was context dependent in the 1 RBD ‘up’ state. While the ‘down’ state RBD in contact with the up state RBD displayed the large shift in position observed in the all ‘down’ state, the down state RBD with its terminal position free displayed an intermediate SD1 configuration. The up state RBD in the u1S2q construct resided largely in the position occupied in the unmutated construct. This indicated the effect of the mutations was primarily isolated to the ‘down’ state and suggested these mutations act to destabilize the ‘down’ state rather than to stabilize the ‘up’ state. These features were largely recapitulated in the u1S2q 2 RBD ‘up’ state conformation with subdomain 1 retaining the shift in the down state RBD (Figure 7). The structural details presented here indicate that, while locking the ‘down’ state RBD into its unmutated position had little impact on the overall configuration of S1, altering the disposition of SD1 had wide ranging impacts, consistent with the observed strain-to-strain differences in the geometric analysis described in Figures 1 and 2.

**Figure 6.**
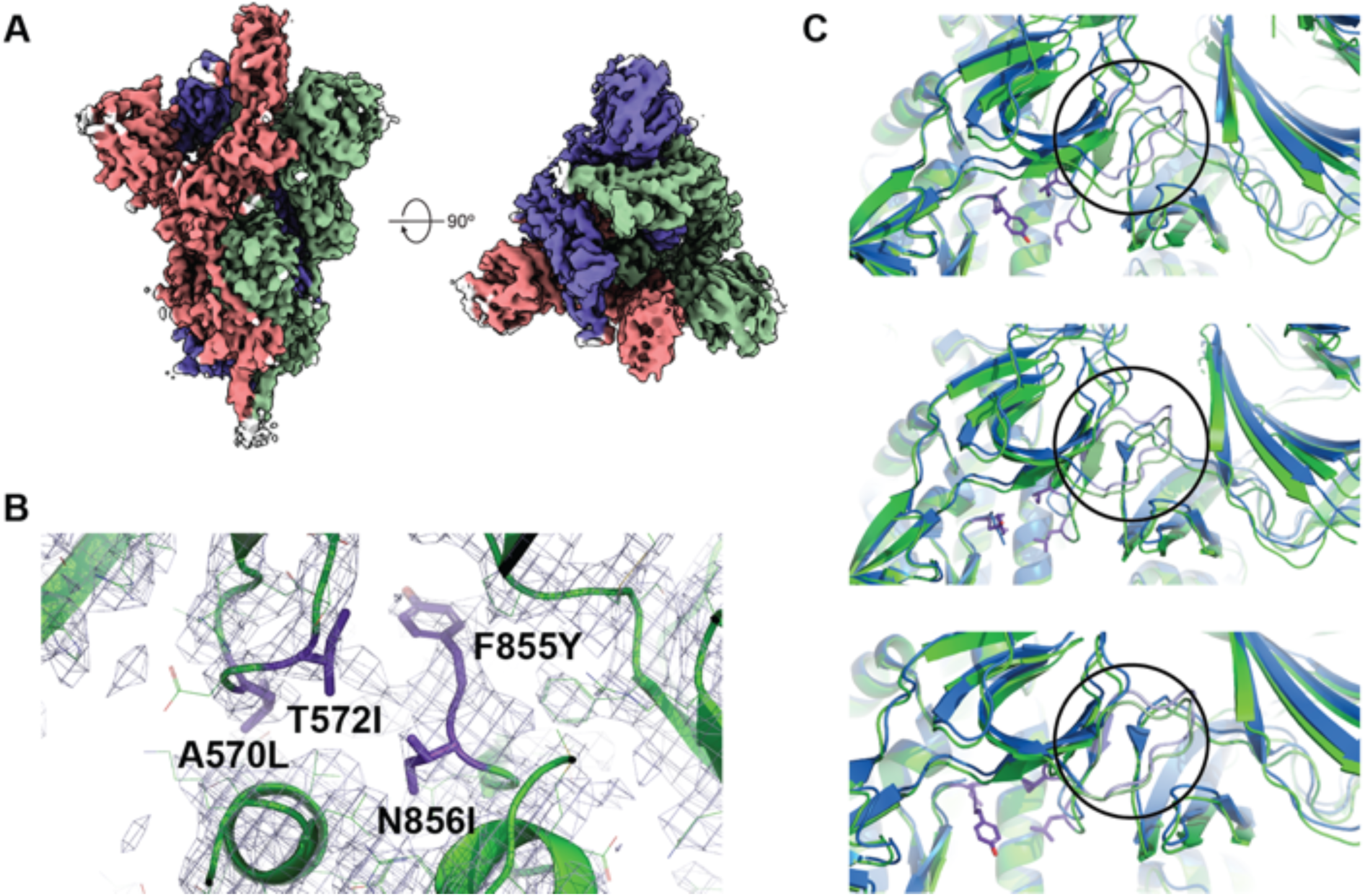
High-resolution structure of the u1S2q 1 RBD ‘up’ state reveals increasing relaxation of the triggered RBDs toward the unmutated structure. **A)** Cryo-EM reconstruction colored by chain. The RBD in the ‘up’ position is marked with an asterisk; side (left) and top (right) views. **B)** Zoomed-in view showing the mutated residues. **C)** (top) Structure of the ‘up’ state RBD coupled ‘down’ state RBD (green) highlighting the shifted subdomain 1 to NTD’ position relative to the unmutated position (blue). (*middle*) Structure of the uncoupled ‘down’ state RBD (green) highlighting the moderately shifted subdomain 1 to NTD’ position relative to the unmutated position (blue). (*bottom*) Structure of the ‘up’ state RBD (green) highlighting the close alignment of subdomain 1 and the NTD’ regions to the unmutated position (blue).

**Figure 7.**
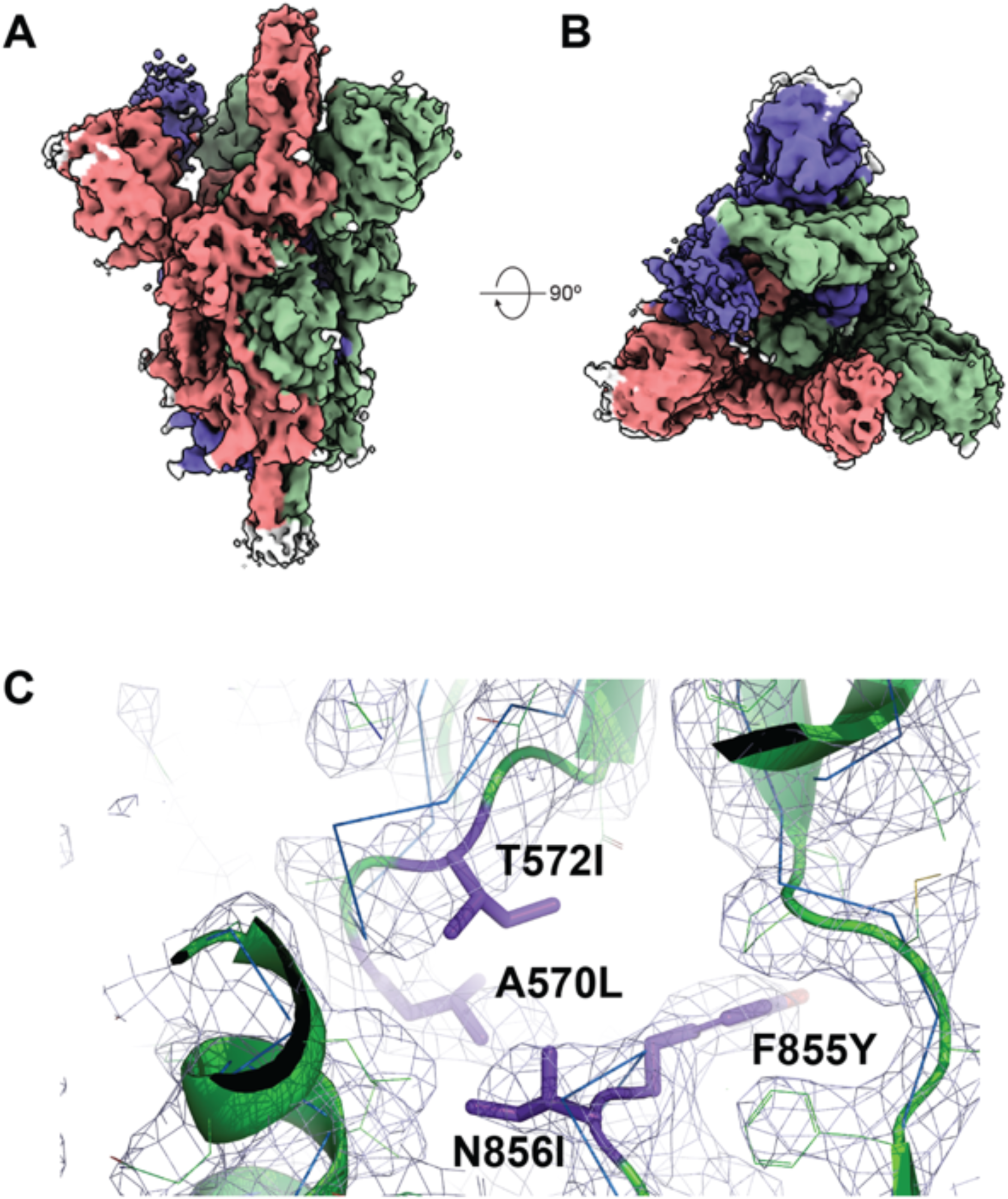
Structure of the u1S2q 2 RBD ‘up’ state indicates modest differences between the 1 RBD ‘up’ state’s subdomain arrangement. **A)** Cryo-EM reconstruction colored by chain. The RBDs in the ‘up’ position are marked with asterisks; side (left) and top (right) views. **B)** Cryo-EM map structural alignment top view. **C)** Structure (green) and cryo-EM map depicting the mutated residue dispositions. The unmutated ‘up’ state protomer alignment is depicted in ribbons (blue).

## Discussion

Conformational plasticity is a hallmark of enveloped-virus fusion-protein structure, owing to the necessity of protecting the conserved viral fusion elements from host immune responses while retaining a sufficiently steep free-energy gradient to enable host cell fusion^16^. Exposed elements are generally well conditioned to be permissive and responsive to mutations through genetic drift and host immune adaptation. Conformational plasticity, however, presents an important difficulty in the context of vaccine and drug design. Indeed, lessons learned in the continued effort to produce a broadly protective HIV-1 vaccine have demonstrated the importance of a detailed understanding and control of fusion protein dynamics^17-28^. The novel SARS-CoV-2 is likely no exception in this regard and indeed the conformational plasticity of the SARS-2 S-protein appeared greater than that of the HIV-1 Env. We aimed to develop a quantitative understanding of β-CoV structural states between strains and within each RBD ‘down’ and ‘up’ state configuration. The wide breadth of domain arrangements along with the relatively small contact area between the S1 and S2 subunits observed here suggested that, despite a relatively low mutation rate, dramatic changes in S-protein structure may occur from few mutations. Indeed, recent evidence for a mutation in the SD2 to S2 contact region suggests a potential fitness gain for acquisition of such interfacial residues^29^. Based upon our results, this mutant, D614G, may indeed alter the conformational landscape of the SARS-CoV-2 S-protein.

From the perspective of immunogen development, the constructs developed here present an opportunity to examine the ability of differentially stabilized S-protein particles to induce two different, yet important antibody responses. First, the disulfide-linked ‘down’ state locked double mutant (rS2d) would presumably eliminate receptor binding site targeting antibodies which make up the majority of observed responses^30,31^. Indeed, a study of MERS responses suggests non-RBD responses (particularly NTD and S2 epitopes) will play an important role in vaccine induced protection^32^. From a theoretical perspective, the wide control over the RBD ‘up’/’down’ distribution available to the virus suggests that, by analogy to known difficult to neutralize HIV-1 strains, conformational blocking of antibody responses would not be unusual. Although this may result in a fitness cost to the virus, it would not necessarily make the virion non-infectious. Using the double mutant rS2d as an immunogen provides a platform from which to induce such non-RBD responses that may be needed to protect against such an evasion. The second area of interest concerns cryptic pocket targeting antibodies which have proven effective in the neutralization of SARS. These antibodies target an epitope presented only in the ‘up’ state RBDs and appear to require a two RBD ‘up’ configuration^33^. The current stabilized ectodomain construct in wide use in SARS-CoV-2 clinical trials was demonstrated previously, and recapitulated here by NSEM, to display only the ‘down’ and one RBD ‘up’ states. However, the u1S2q, SD1/S2 targeting design developed here display a prominent two RBD ‘up’ state distribution compatible with these cryptic-epitope targeting MAbs. This suggests it is capable of inducing such antibodies. While complicating factors, such as a potential for vaccine enhancement, may favor the use of truncated, single domain constructs which may display fewer potentially weakly or non-neutralizing epitopes, these, along with the designs presented here will allow for a detailed characterization of not only vaccine immunogenicity but also antigenicity, paving the way for next generation vaccines for the novel SARS-CoV-2 and the eventual development of a broadly neutralizing β-CoV vaccine. Thus, while the previous generation of stabilizing mutations ensure well folded trimer, the rational design approach developed here provides a means by which precisely controlling the RBD orientation distribution, thus allowing exploratory efforts to understand the role of conformational dynamics from the perspective of vaccine and drug development.

## Acknowledgements

Initial cryo-EM data were collected on the Titan Krios at the Shared Materials and Instrumentation Facility in Duke University. Data collection for high-resolution structure determination was performed at the National Center for Cryo-EM Access and Training (NCCAT) and the Simons Electron Microscopy Center located at the New York Structural Biology Center, supported by the NIH Common Fund Transformative High Resolution Cryo-Electron Microscopy program (U24 GM129539,) and by grants from the Simons Foundation (SF349247) and NY State. We thank Ed Eng, Dajia Dobe, Mark Walters and Holly Leddy for microscope alignments and assistance with cryo-EM data collection as well as Jason McLellan for providing the plasmid for the unmutated construct. This work was supported by UM1 AI100645 (B.F.H), the Duke Center for HIV/AIDS Vaccine Immunology-Immunogen Discovery, and UM1 AI44371 (B.F.H.), the Duke Consortium for HIV/AIDS Vaccine Development, Division of AIDS, NIAID, NIH; Duke University Center for AIDS Research (CFAR); Translating Duke Health Initiative (P.A. and R.C.H.), and the Intramural Research Program of the NIH; National Institute of Environmental Health Sciences (ZIC ES103326 to M.J.B.).

## Author contributions

R.C.H and P.A. conceived the study, determined cryo-EM structures and analyzed data. R.C.H. led computational studies. P.A. led structural studies. R.J.E led NSEM studies and analyzed data. K.M. collected NSEM data and performed initial data analysis. K.J. and V.S. produced and purified proteins. S.G. purified proteins. M.K. and A.H. optimized cryo-EM specimen. K.J. and R.P. performed binding studies to optimize spike preparations. R.J.E., M.B., B.F.H and P.A. supervised studies. R.C.H and P.A. wrote the manuscript with help from all authors.

## Data Availability

The datasets generated and/or analyzed during this study are available from the corresponding authors on reasonable request.

## Code Availability

The code developed during this study is available from the corresponding author on reasonable request.

## Methods

### Vector based analysis

Vector analysis was performed using available cryo-EM structures for SARS-2^13,14^, SARS^4,5,7,8^, MERS^4,12^, and other human^2,10^ and murine^11^ β-CoV spike proteins. Domains for the vector analysis were selected based upon visual inspection of alignments between SARS, MERS, and SARS-CoV-2 structures. Specifically, C_α_ centroids for the S1 NTD, RBD, SD1, SD2 (SARS-CoV-2 residues, 27-43 and 54-271, 330-443 and 503-528, 323-329 and 529-590, 294-322 and 591-696, respectively; equivalent SARS/MERS/Murine/HKU1/OC43 residues selected based upon structural alignment with SARS-CoV-2) as well as a β-sheet motif in the NTD (residues 116-129 and 169-172) and a helix motif in the RBD (residues 403-410) were determined. The NTD was split into two regions with the SD1 contacting, SD2 adjacent portion referred to here as the NTD’ (residues 44-53 and 272-293). C_α_ centroids in the S2 domain were obtained for a β-sheet motif (residues 717-727 and 1047-1071) and the CD domain (711-716 and 1072-1122). Vector magnitudes, angles, and dihedrals between these centroids were determined and used in the subsequent analysis. Vector analysis was performed using the VMD^34^ Tcl interface. Principal component analysis performed in R with the vector data centered and scaled^35^.

### Rational, structure-based design

Structures for SARS (PDB ID 5X58^4^), MERS (PDB ID 6Q04^36^), and SARS-CoV-2 (PDB ID 6VXX^15^) were prepared in Maestro^37^ using the protein preparation wizard^38^ followed by *in silico* mutagenesis using Schrödinger’s cysteine mutation^39^ and residue scanning^40^ tools. Residue scanning was first performed for individual selected sites allowing mutations to Leu, Ile, Trp, Tyr, and Val followed by scanning of combinations for those which yielded a negative overall score. Scores and visual inspection were used in the selection of the prepared constructs.

### Protein expression and purification

The SARS-CoV-2 ectodomain constructs were produced and purified as described previously (ref-Wrapp et al., McLellan). Briefly, a gene encoding residues 1−1208 of the SARS-CoV-2 S (GenBank: MN908947) with proline substitutions at residues 986 and 987, a “GSAS” substitution at the furin cleavage site (residues 682–685), a C-terminal T4 fibritin trimerization motif, an HRV3C protease cleavage site, a TwinStrepTag and an 8XHisTag was synthesized and cloned into the mammalian expression vector pαH. All mutants were introduced in this background. Expression plasmids encoding the ectodomain sequence were used to transiently transfect FreeStyle293F cells using Turbo293 (SpeedBiosystems). Protein was purified on the sixth day post-transfection from the filtered supernatant using StrepTactin resin (IBA).

### Cryo-EM sample preparation and data collection

Purified SARS-CoV-2 spike preparations were diluted to a concentration of ∼1 mg/mL in 2 mM Tris pH 8.0, 200 mM NaCl and 0.02% NaN3. 2.5 µL of protein was deposited on a CF-1.2/1.3 grid that had been glow discharged for 30 seconds in a PELCO easiGlow™ Glow Discharge Cleaning System. After a 30 s incubation in >95% humidity, excess protein was blotted away for 2.5 seconds before being plunge frozen into liquid ethane using a Leica EM GP2 plunge freezer (Leica Microsystems). Frozen grids were imaged in a Titan Krios (Thermo Fisher) equipped with a K3 detector (Gatan). Data were acquired using the Leginon system^41^. The dose was fractionated over 50 raw frames and collected at 50ms framerate. This dataset was energy-filtered with a slit width of 30 eV. Individual frames were aligned and dose-weighted. CTF estimation, particle picking, 2D classifications, *ab initio* model generation, heterogeneous refinements, and homogeneous 3D refinements were carried out in cryoSPARC^42^.

### Cryo-EM structure fitting

Structures of the all ‘down’ state (PDB ID 6VXX) and single RBD ‘up’ state (PDB ID 6VYB) from the previously published SARS-CoV-2 ectodomain were used to fit the cryo-EM maps in Chimera^43^. The 2 RBD ‘up’ state was generated in PyMol using the single RBD ‘up’ state structure. Mutations were made in PyMol^44^. Coordinates were then fit manually in Coot^45^ followed by iterative refinement using Phenix^46^ real space refinement and subsequent manual coordinate fitting in Coot. Structure and map analysis were performed using PyMol and Chimera.

---

**Supplemental Figure S1.**
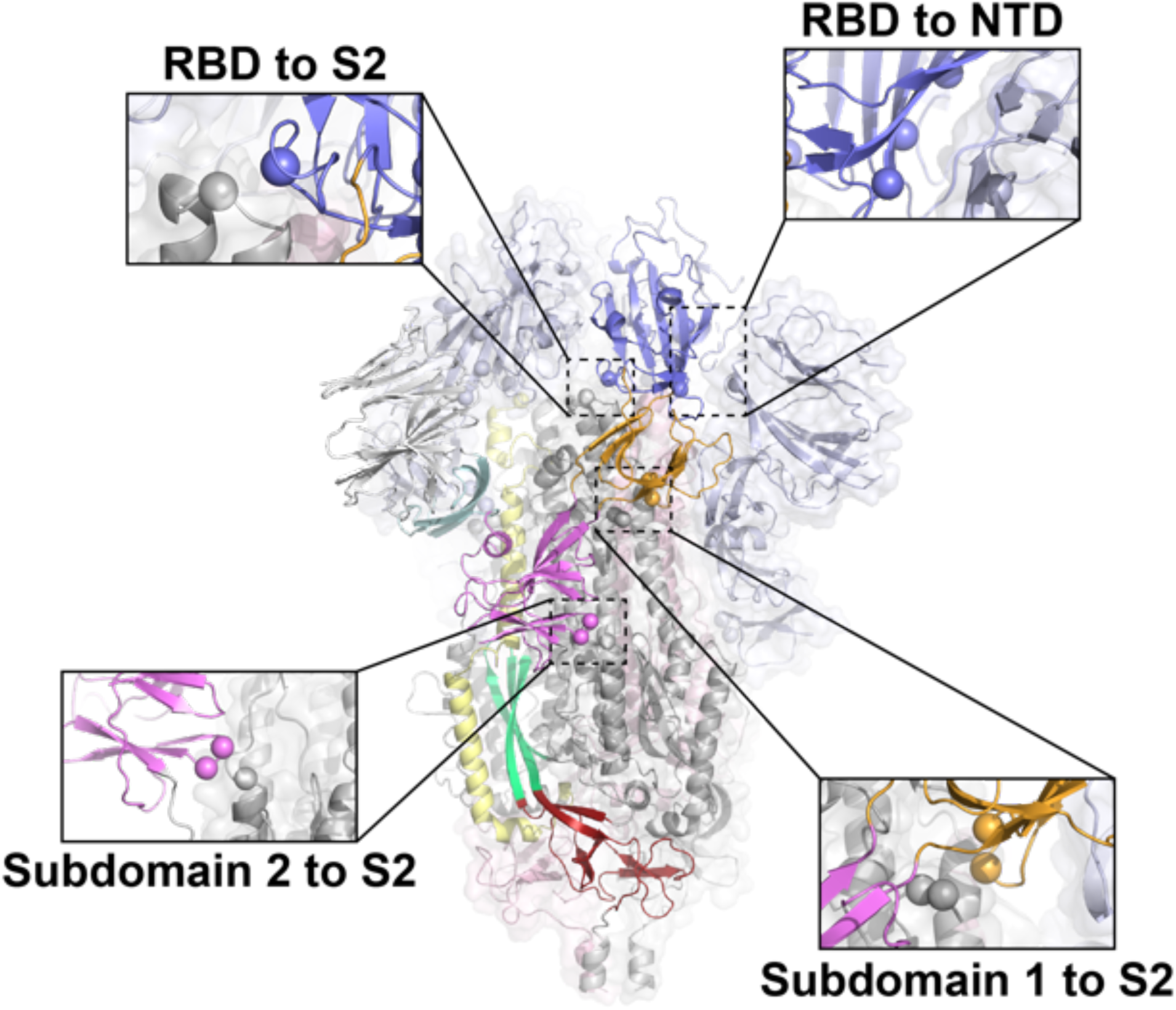
Sites identified for differential stabilization of the SARS-CoV-2 S-protein. Single protomer colored according to Figure 1 with remaining two protomers color according to S1 (light blue) and S2 (grey). Spheres indicate candidate mutation sites.

**Figure S2.**
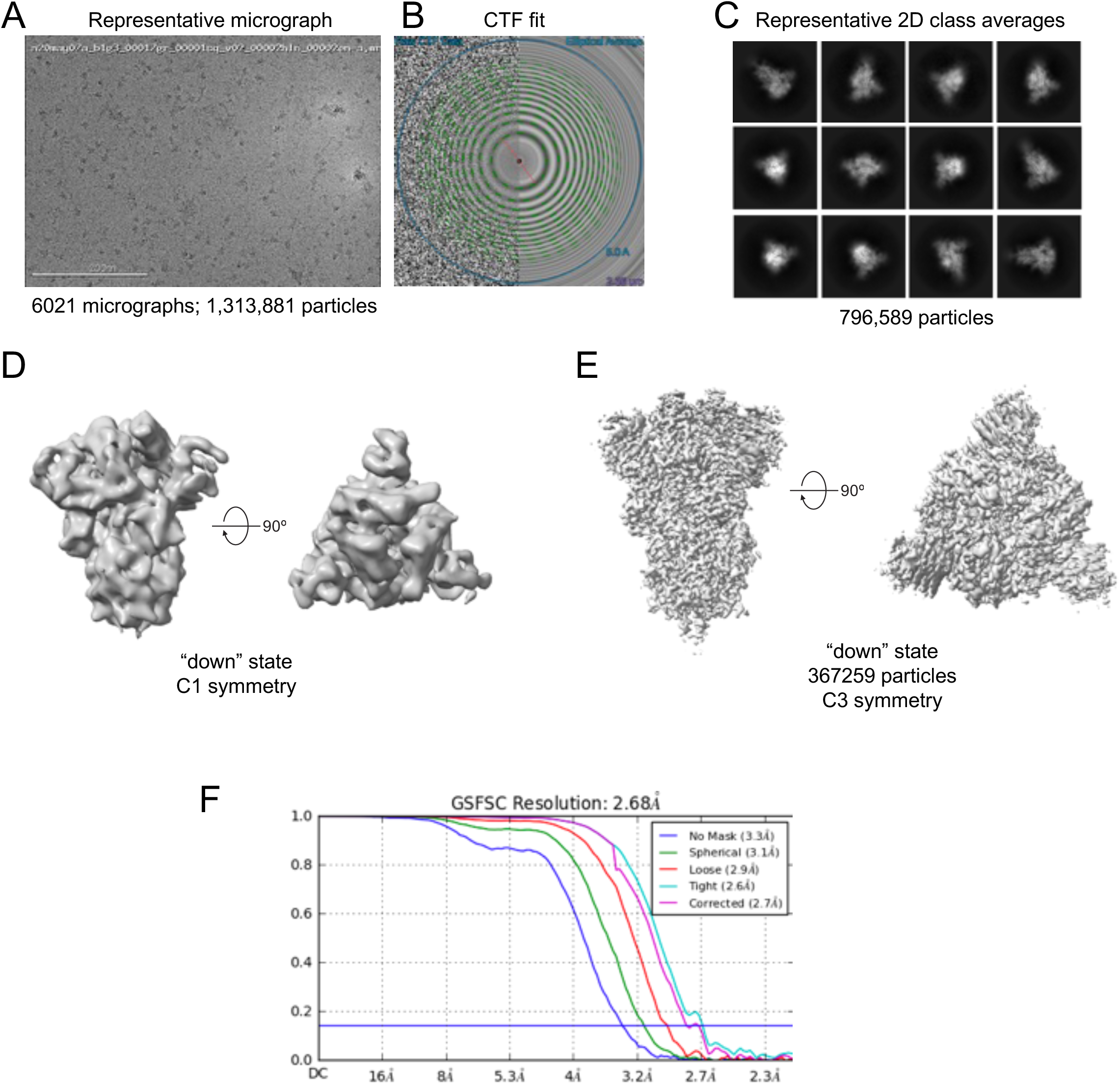
Cryo-EM data processing details for r2S2d. **A)** Representative micrograph. **B)** CTF fit **C)** Representative 2D class averages. **D)** Ab initio reconstruction. **E)** Refined map. **F)** Fourier shell correlation curves.

**Figure S3.**
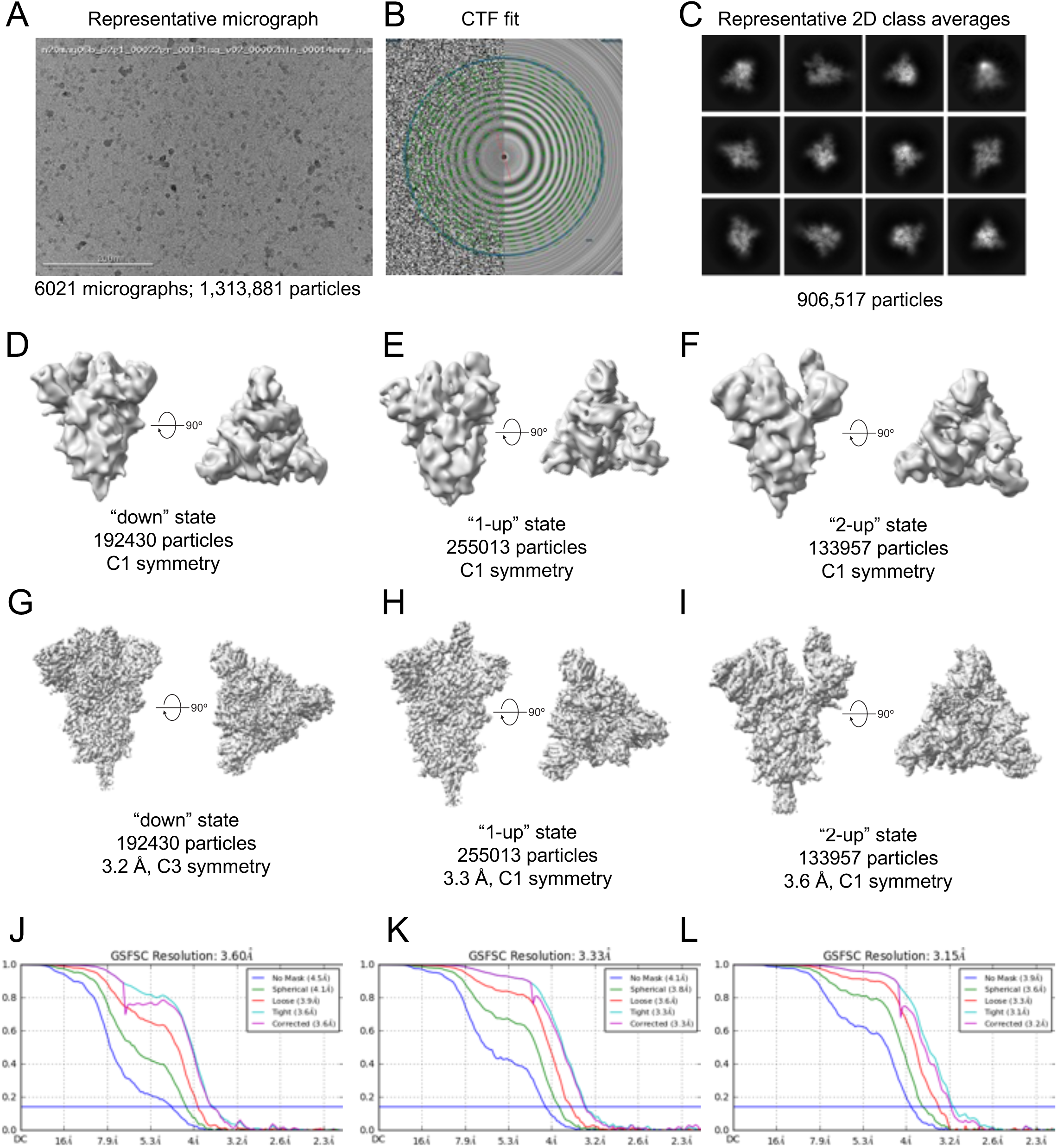
Cryo-EM data processing details for u1s2q. (A) Representative micrograph. (B) CTF fit (C) Representative 2D class averages. (D-F) *Ab initio* reconstructions for the (D) “down” state, (E) “1-up” state and (F) “2-up” state. (G-I) Refined maps for the (G) “down” state, (H) “1-up” state and (I) “2-up” state. (J-L) Fourier shell correlation curves for the (J) “down” state, (E) “1-up” state and (F) “2-up” state.

**Supplemental Figure S4.**
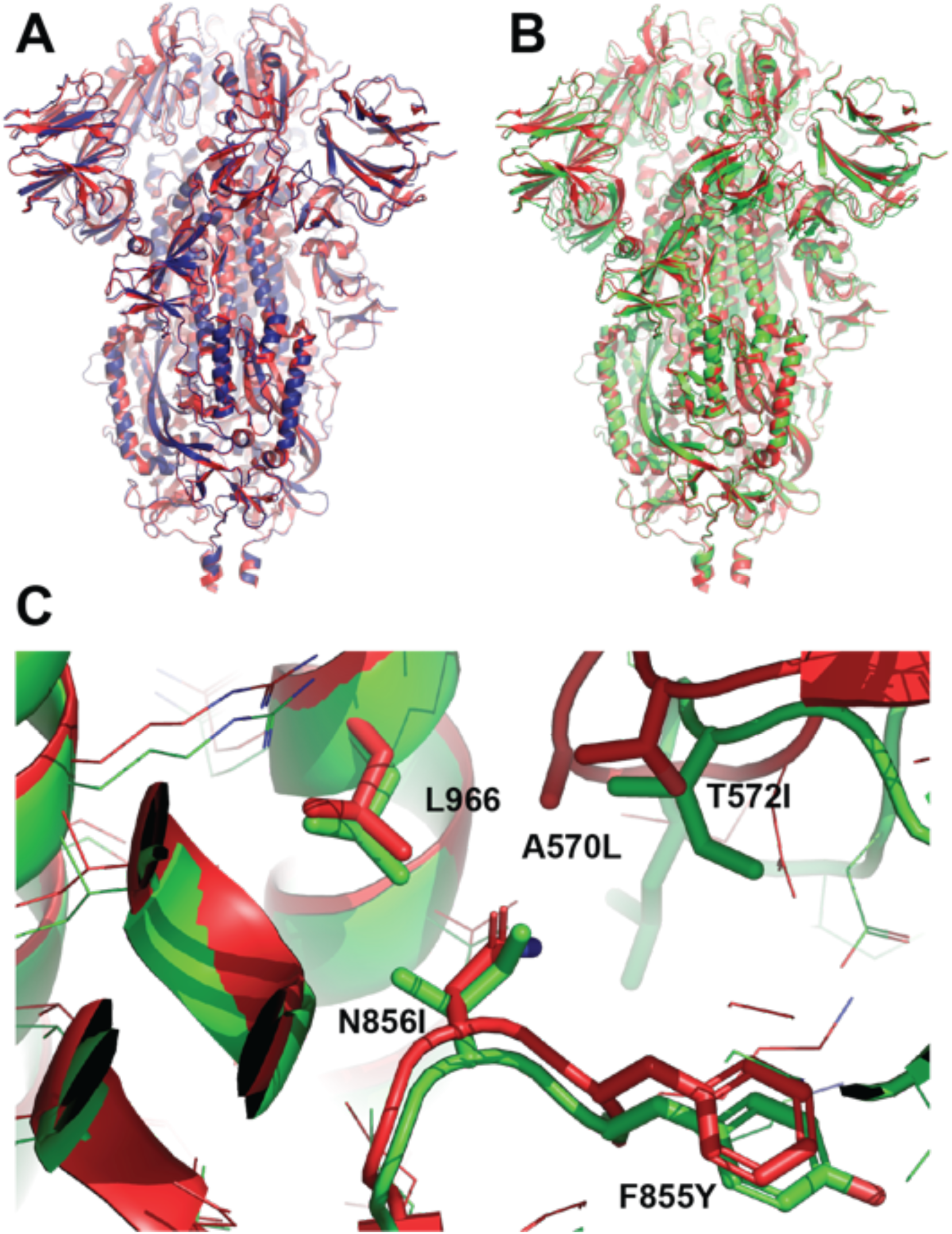
Alignment of the rS2d and u1S2q designs with the unmutated construct. **A)** Structure of rS2d (dark blue) aligned to the unmutated construct (PDB ID 6VXX; red). **B)** Structure of u1S2q (green) aligned to the unmutated construct (PDB ID 6VXX; red). **C)** The u1S2q (green) mutation sites compared to the unmutated form (red).

## Notes

### Competing Interest Statement

The authors have declared no competing interest.

